# UBE3A reinstatement restores behavior and proteome in an Angelman Syndrome mouse model of Imprinting Defects

**DOI:** 10.1101/2024.09.29.615689

**Authors:** Claudia Milazzo, Ramanathan Narayanan, Solveig Badillo, Silvia Wang, Rosaisela Almand, Edwin Mientjes, Stormy Chamberlain, Thomas Kremer, Ype Elgersma

**Affiliations:** Dept. of Clinical Genetics; Erasmus MC Center of Expertise for Neurodevelopmental Disorders (ENCORE); Erasmus MC, Rotterdam, The Netherlands; Neuroscience and Rare Diseases (NRD), Pharmaceutical Research & Early Development, F. Hoffmann-La Roche Ltd., Basel, Switzerland; Predictive Modelling and Data Analytics (PMDA), Pharmaceutical Sciences, F. Hoffmann-La Roche Ltd., Basel, Switzerland; Biomarker Development – Translational Medicine, Novartis Institutes for Biomedical Research, Novartis Pharma Ltd., Basel, Switzerland

**Author notes:** Address correspondence to: Ype Elgersma, Department of Clinical Genetics, Erasmus MC, 3015 GD Rotterdam, the Netherlands. Contributed equally to this work (Claudia Milazzo, Ramanathan Narayanan). Shared corresponding author (Stormy Chamberlain, Thomas Kremer, Ype Elgersma). **Conflict of interest:** This work was partly funded by F. Hoffmann-La Roche Ltd.

## Abstract

Angelman Syndrome (AS) is a severe neurodevelopmental disorder wionly symptomatic treatment currently available. Besides mutations within the *UBE3A* gene, AS is caused by deletions, imprinting center defects (mICD) or uniparental disomy of chromosome 15 (UPD). Current mouse models are *Ube3a*-centric and do not address expression changes of other 15q11-q13 genes on AS pathophysiology. Here, we studied a mouse line that harbors a mutation affecting the AS-PWS imprinting center, hence modeling mICD/UPD AS subtypes. mICD mice showed significant reduction in UBE3A protein, bi-allelic expression of *Ube3a-ATS* and *Mkrn3-Snord115* gene cluster, leading to robust AS behavioral deficits and proteome alterations similar to *Ube3aKO* mice. Genetic UBE3A overexpression in mICD mice, mimicking therapeutic strategies that effectively activate the biallelic silenced *Ube3a* gene, resulted in a complete rescue of all behavioral and proteome alterations. Subsequently, treatment with an antisense oligonucleotide (ASO) to directly activate the biallelic silenced *Ube3a* gene in mICD mice also resulted in efficient reinstatement of UBE3A, alongside a partial rescue of behavioral phenotypes. Taken together, these findings demonstrate that UBE3A loss is the primary factor underlying AS phenotypes in the mICD/UPD mouse model, and also corroborate that UBE3A reinstatement is an attractive therapeutic strategy for mICD/UPD AS individuals.

## Introduction

Angelman Syndrome (AS) is a severe debilitating neurodevelopmental disorder, with an estimated incidence of 1 in 20,000 (1). AS manifests with strong deficits in fine and gross motor skills, absence of speech, intellectual disability, and aberrant behavior (2). Additionally, 80% of the patients have epilepsy and sleep disturbances (2, 3). Currently, available treatments focus on symptom management, particularly seizure reduction and sleep improvement. AS is caused by the loss of functional maternally expressed UBE3A protein in neurons (4, 5). *UBE3A* encodes an E3 ubiquitin ligase involved in protein degradation and transcriptional co-activation, critical for proper neurodevelopment and synaptic function (reviewed by (6). The genetic etiology underlying loss of UBE3A is diverse and varies from large genomic deletions of the 15q11-q13 region in about 60-70% of AS cases, imprinting defects affecting the AS-imprinting center (ICD) or paternal uniparental disomy of chromosome 15 (UPD), in about 20% of AS individuals, as well as single nucleotide mutations specifically affecting the *UBE3A* gene, affecting approximately 15% of AS patients (3, 4, 7).

Since all aforementioned mutations affect UBE3A expression in the brain, AS mouse studies have primarily focused on the *Ube3a* mouse model. Recent studies in *Ube3a* animal models have demonstrated the potential of UBE3A reinstatement via antisense oligonucleotide (ASO) treatment or gene therapy as a disease-modifying strategy for AS (8–11). These ASOs target the *UBE3A-ATS*, a long non-coding transcript encoded by the distal end of *SNHG14*, which is responsible for in-cis silencing of the paternal *Ube3a* allele (12–15) (Figure S1A). However, these *Ube3a-*centric approaches do not address the consequence of expression changes of other genes in the 15q11-q13 locus in individuals with large genomic deletions, UPD or mICD mutations. More specifically, UPD and mICD mutations not only result in loss of expression of the maternal *UBE3A* gene, but also cause biallelic expression of genes in the 15q11-q13 region, normally expressed only from the paternal allele, (i.e. *MKRN3, NDN, MAGEL2* genes located upstream of the AS- IC) as well as biallelic expression of the *SNHG14* polycistronic transcription unit including the *UBE3A-ATS* (13, 16). Notably, the biallelic expression of *UBE3A-ATS*, poses risks associated with UBE3A protein over- dosage upon ASO-mediated targeting of the *UBE3A-ATS* to achieve *UBE3A* reactivation (Figure S1A). For example, chromosomal duplications of the 15q11-13 region or gain-of-function mutations that result in increased UBE3A levels or activity are associated with developmental delay, neuropsychiatric disorders, and autism spectrum disorder (ASD) (reviewed by (17)). However, overexpression of UBE3A in mice appeared largely insufficient to cause discernable phenotypes, suggesting that overexpression of UBE3A is only detrimental in combination with overexpression of other genes in the 15q11-13 locus (18).

In this study we made use of a mICD mouse model to address the effect of increased expression of genes located upstream of the AS-IC, and we investigated the effect of expressing two *Ube3a* gene copies in this model. We found that the behavioral deficits and proteomic alterations observed in mICD mice were completely reversed upon UBE3A overexpression. Treatment with ASO resulted in effective reinstatement of UBE3A and a partial behavioral rescue. Taken together, these results indicate that UBE3A is causal of AS pathophysiology in mICD and UPD, while other deregulated genes play an undetectable role in this experimental setting. In light of these findings, we believe that mICD and UPD patients could benefit from UBE3A reinstatement therapies currently tested in clinical trials using ASOs.

## Results

### mICD mice show robust anatomical and behavioral AS phenotypes

Elegant studies have shown that the insertion of a transcription termination cassette between the AS and PWS imprinting centers disrupt imprinting in mice (depicted in Figure S1A), which makes this mouse line (Rr70^em1Rsnk^; MGI: 6306477) a very good model with high construct validity for studying the mICD and UPD AS subtypes (19).

We used this line (in this study referred to as ‘mICD’ mice) and control littermates in the F1 hybrid 129S2-C57BL/6J background to assess previously reported gene expression changes of the *Ube3a-ATS, Mkrn3, Snrpn, Snord 115,* and *Snord 116* genes. We observed a two-fold increase in gene expression of these genes, in line with previous findings. The *Ube3a* transcript and protein were reduced by approximately 80% relative to WT controls in the cortex of adult mICD mice (Figure S1 B-H, Table S2). To investigate whether mICD mice show robust AS behavioral features, we made use of a behavioral test battery well established in *Ube3a*-centric mouse lines that revealed motor impairment, increased anxiety and deficiencies in species-specific innate behaviors (Figure 1A) (20). Adult mICD mice and control littermates in the F1 hybrid 129S2-C57BL/6J background were used for the behavioral battery and following analyses. Similar to *Ube3a* mice (11, 20), mICD mice showed marked behavioral deficiencies in the reverse rotarod test, hind-limb clasping, marble burying, and in the nest building task. In addition, immobility in the tail suspension test was increased (Figure 1 A-H; Figure S2; Table S2). Similar to mice lacking UBE3A, mICD mutants showed increased body weight and microcephaly compared to WT mice (Figure 1I and J) (20, 21). In contrast, we did not find a difference in distance moved by mICD mice compared to WT controls in the open field test (Figure 1C).

**Figure 1.**
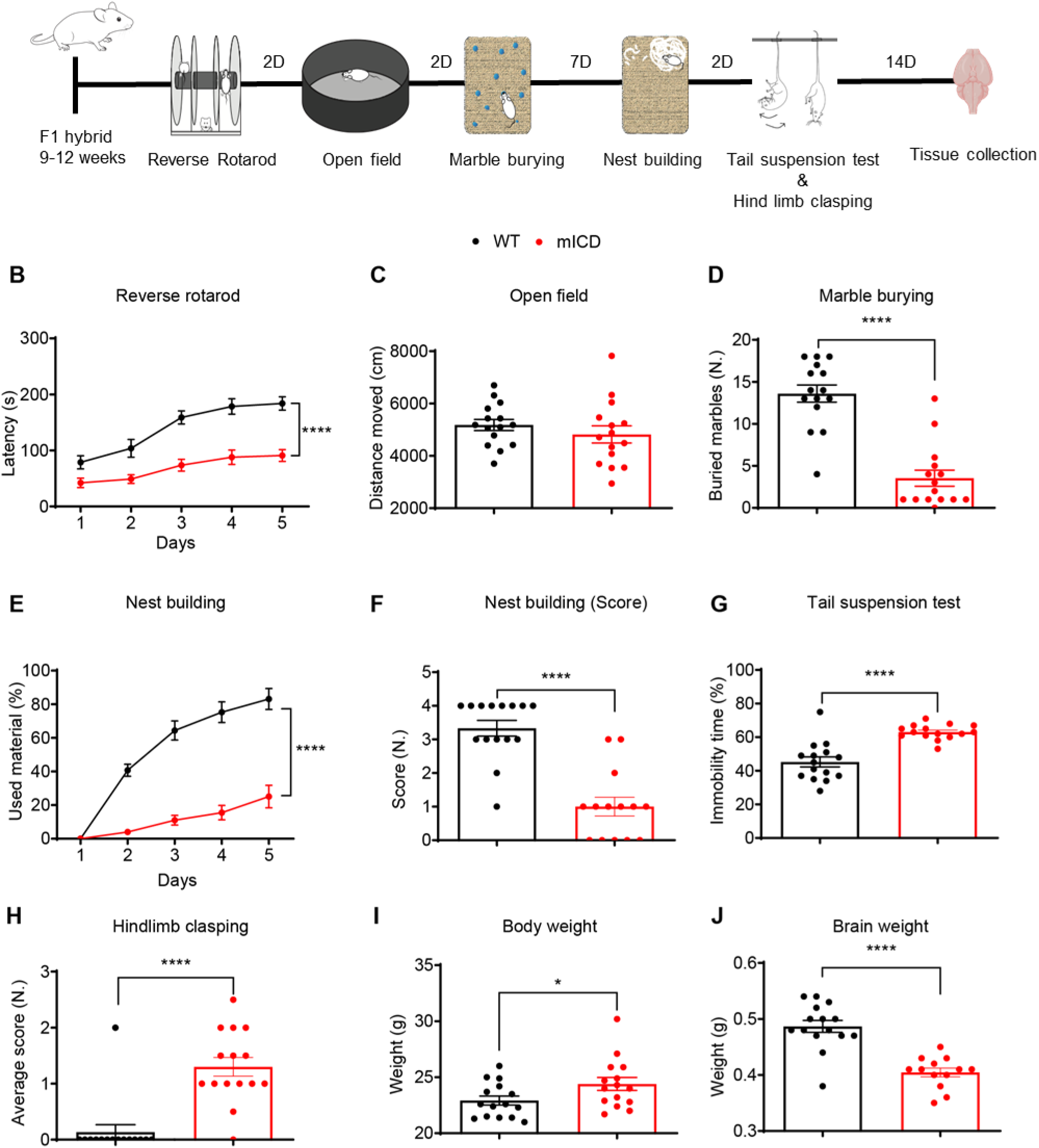
mICD mice show robust anatomical and behavioral AS phenotypes. **A.** Illustration of the timeline for execution of the behavioral battery and tissue collection. (B-H) Behavioral battery performed with WT (black) and mICD (red) mice in the reverse rotarod (n=15; 15), open field (n= 15, 15), marble burying (n= 15, 15), nest building (n= 15, 15), quality of built nest (n= 15, 14), tail suspension (n= 15, 15) and hind-limb clasping (n= 15, 15) tests. (I-J) Anatomical features such as body (n= 15, 15) and brain weight (n= 15, 13) were measured at the beginning and the end of the battery, respectively. A T-test or 2-way ANOVA was performed for statistical analysis. Data are represented as means ± SEM. *P* values are displayed as asterisks in the figure: not shown if *P* > 0.05, \**P* ≤ 0.05, \*\**P* ≤ 0.01, \*\*\**P* ≤ 0.001, \*\*\*\**P* ≤ 0.0001.

### Proteomics reveals broader protein dysregulation in mICD mice compared to *Ube3a KO* mice, but with similar top hits

Several studies have shown that UBE3A acts both as a coactivator of nuclear/steroid hormone receptors (22–25) and as a E3 ubiquitin ligase that impacts proteasomal, tRNA synthetase and synaptic pathways (26). In addition, changes in such key cellular pathways can in turn lead to secondary and tertiary effects on transcripts and proteins. To identify quantitative molecular phenotypes in mICD mice, we performed total RNA sequencing and data independent acquisition (DIA) -based proteomics.

Total RNA sequencing of cortex tissue from adult mICD and WT controls showed significant downregulation of *Ube3a*, increased expression of *Ube3a-ATS*, *Snrpn* and *Ndn*, confirming our RT-qPCR findings (Figure 2A, B). In addition, several transcripts that map to the same chromosomal region also show increased expression (highlighted in blue). Besides these major genotype-driven changes, there is only a modest dysregulation of transcripts (0.75 < logFC < -0.75, n= 22 downregulated, n= 36 upregulated) without an enrichment for specific pathways. Notable examples are (with logFC values): *Adora2a* (2.7), *Rgs9* (1.9), *Oprk1* (1.8), which are upregulated in mICD and *Mdfic2* (-1.8), *Prl* (-2.4), *Gh* (-2.6), which are downregulated in mICD (Table S3).

**Figure 2.**
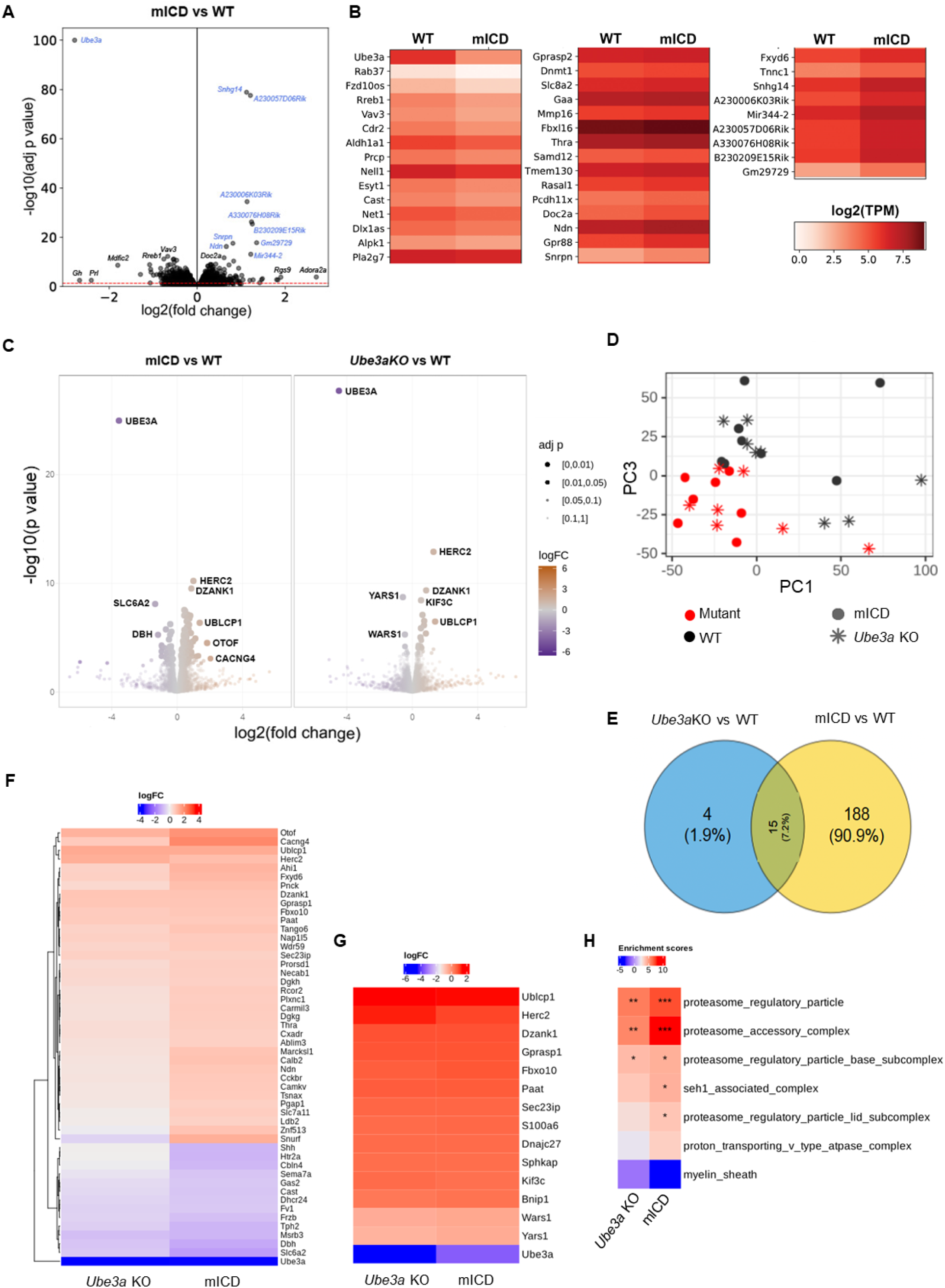
Proteomics reveals broader protein dysregulation in mICD compared to *Ube3a KO* mice, with similar top hits. (A) Volcano plot of –log10(adjusted p-value) vs log2(fold change) of total RNA sequencing comparing cortex of adult mICD and WT mice. Transcripts originating from the 15q11-q13 locus on mouse chromosome 7 are highlighted in blue. Red line corresponds to p-value of 0.05 (B) Heatmap of the total RNAseq data showing top 40 downregulated (left panel) and upregulated (center and right panels) transcripts depicted as log2TPM, filtered by adjusted p-value (C) Volcano plot showing the magnitude of effect (log2 fold change on the x-axis) against the statistical significance (-log10 FDR-adjusted *p* value on the y-axis) for the data independent acquisition (DIA)-based proteomic profiling of mICD (n = 8) and *Ube3a KO (*n= 8) cortex (with respect to their WT controls, n = 8 each) (D) Principal component analysis (PCA) of all proteins shows a clear separation based on genotype (PC 3) and a minimal separation based on mouse model (PC 1). (E) Venn diagram showing an overlap of significant hits between the two mouse models (FDR<0.05). (F) Heatmap showing the top 50 differentially abundant proteins in the two mouse models for the contrasts mICD vs WT and *Ube3a KO* vs WT (red: most abundant; blue: least abundant). Hierarchical clustering was used to group proteins based on FDR. (G) Heatmap of the 15 proteins that are commonly dysregulated. (H) Significantly enriched gene sets from the Gene Ontology - Cellular Component (GO:CC) are shown for the contrasts mICD vs WT and *Ube3a KO* vs WT (*** Benjamini Hoch FDR<0.001, **FDR<0.01, *FDR<0.05).

DIA proteomics of cortex tissue from adult mICD mice was performed together with *Ube3a^p+/m–^* mice (from here on referred to as *‘Ube3a KO’* mice) for comparability. The mICD mouse cortex showed dysregulation of more proteins (n= 145 upregulated, n= 58 downregulated, FDR<0.05) than that of *Ube3a KO* mice (n= 15 upregulated, n= 4 downregulated) (both with respect to their WT controls) (Figure 2C, Table S3). Principal component analysis of all proteins (PCA) separated the samples by genotype (PC 3), whereas only a minimal separation is observed based on the mouse model (PC 1) (Figure 2D). This is reflected in a strong overlap between the two mice models, with identical top protein hits (Figure 2E, F). The 14 commonly dysregulated proteins (excluding UBE3A) are HERC2, DZANK1, KIF3C, FBXO10, SEC23IP, PAAT, S100A6, UBLCP1, GPRASP1, SPHKAP, BNIP1, DNAJC27, which are upregulated and YARS1 and WARS1, which are downregulated (Figure 2G). Notably, geneset enrichment analysis using the cellular component geneset (GO:CC) showed significant enrichment of proteasome related terms both in the mICD and *Ube3a KO* mice (Figure 2H, Table S3).

### Transgenic UBE3A overexpression in mICD mice rescues behavioral and proteomic alterations

To evaluate the role of the bi-allelically expressed paternal genes on AS pathophysiology and to assess whether bi-allelic expression of *Ube3a* in neurons is safe and effective to reverse the identified phenotypes, we crossed mICD mice with *Ube3a^OE^* mice for comprehensive molecular and behavioral characterization. The *Ube3a^OE^* line harbors two copies of a *Ube3a* containing bacterial artificial chromosome (BAC) in chromosome 3 as previously described (18). Quantitative PCR and western blot analysis indicate bi-allelic expression of UBE3A in multiple brain regions of mICD-*Ube3a^OE^* mice, with a 2.2, 1.6 and 1.9 fold increase of UBE3A protein compared to WT controls in the cortex, hippocampus and striatum respectively (Figure 3A, B) (18).

**Figure 3.**
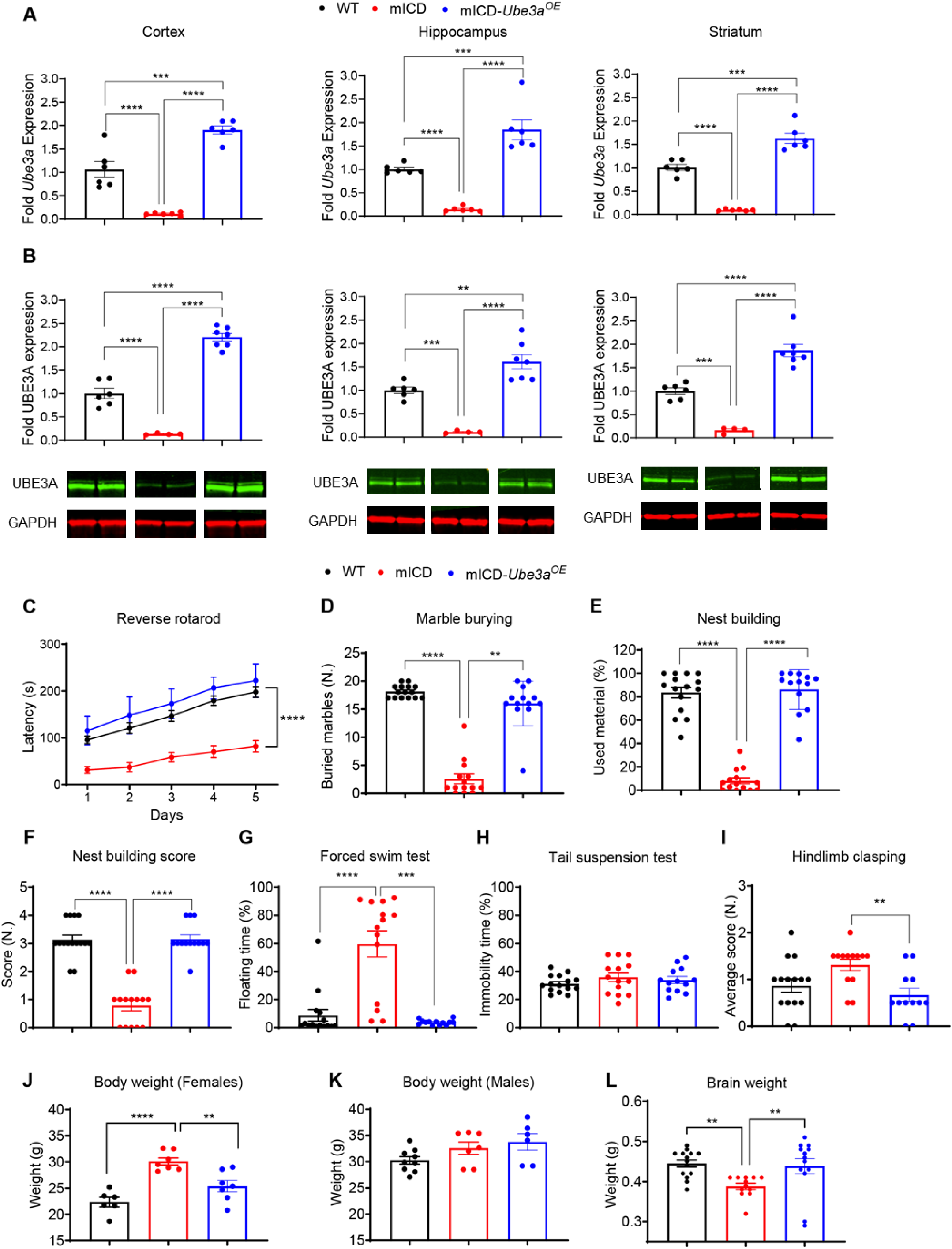
Embryonic transgenic UBE3A reinstatement in mICD mice rescues all behavioral deficits. (A) Quantitative RT-PCR analyzing *Ube3a* mRNA in cortex, hippocampus and striatum lysates of WT, mICD and mICD-*Ube3a^OE^* (n=6 each) and (B) Western blot analyzing UBE3A protein in cortex, hippocampus and striatum lysates of WT (n = 6), mICD (n = 4), and mICD-*Ube3a^OE^* mice (n = 7), obtained 1 week after completion of the behavioral battery. Two bands representing the nuclear and cytosolic isoforms of UBE3A, can be detected at 100 kDa and GAPDH, here used as loading control, at 37 kDa. (C-I) Behavioral battery performed with WT (black), mICD (red) and mICD-*Ube3a^OE^* (blue) mice in the reverse rotarod (n = 15, 14, 13), marble burying (n = 15, 14, 13), nest building (n = 15, 14, 13), quality of built nest (n = 15, 14, 13), forced swim (n = 15, 14, 13), tail suspension (n = 15, 14, 13) and hind-limb clasping tests (n = 15, 14, 12). (J-L) Anatomical features such as body (Females n= 6, 7, 7; Males n = 9, 7, 6) and brain weight (n = 15, 14, 13) were measured at the beginning and end of the battery, respectively. A One-way ANOVA or 2-way ANOVA was performed for statistical analysis. Data are represented as means ± SEM. *P* values are displayed as asterisks in the figure: not shown if *P* > 0.05, \**P* ≤ 0.05, \*\**P* ≤ 0.01, \*\*\**P* ≤ 0.001, \*\*\*\**P* ≤ 0.0001.

We hypothesized that embryonic reinstatement of UBE3A in the transgenic mICD-*Ube3a^OE^* should prevent the manifestation of all characteristic AS phenotypes, unless the paternally expressed genes do have a significant role on AS pathophysiology, or if there is an interaction between the paternally expressed genes and increased UBE3A levels. Indeed, we observed a rescue of all behavioral deficits of the mICD mice upon overexpression of UBE3A (Figure 3C-H, Figure S3A, Table S2). However, the tail suspension phenotype and hindlimb clasping deficits observed in the initial mICD cohort, were not replicated in this new crossing.

Regarding typical AS anatomical features, female mICD mice showed a significant increase in body weight compared to WT mice, which was absent in mICD- *Ube3a^OE^* mice. No significant difference in body weight was observed among males (Figure 3 J, K). Interestingly, while mICD and WT mice maintained a similar body weight from the start to the end of the behavioral battery, the body weight of both male and female mICD- *Ube3a^OE^* mice showed a noticeable increase over time (Figure S3B, Table S2). Also the microcephaly observed in the mICD mice was fully rescued in the mICD-*Ube3a^OE^* mice (Figure 3L).

To assess the impact of UBE3A overexpression in the mICD mice at the proteome level, DIA proteomics of cortex tissue from adult mICD-*Ube3a^OE^*, mICD and WT control mice were performed (Figure 4A, Table S3). Principal component analysis of all proteins (PCA) separated the samples by both genotype (PC 2) and condition (PC 1), wherein the mICD-*Ube3a^OE^* proteome is closer to WT than mICD proteome (Figure 4B). The top 50 significant protein hits in the mICD cortex were either fully or partially normalized, or in many cases completely reversed in the mICD-*Ube3a^OE^* cortex (Figure 4C). In addition, the 14 proteins identified as commonly differentially expressed between mICD and *Ube3a-KO* mice, were largely normalized, but not reversed (Figure 4D). Similarly, geneset enrichment analysis using the cellular component geneset (GO:CC) showed a clear rescue of pathway alterations, particularly proteasome related, in the mICD-*Ube3a^OE^* cortex (Figure 4E, Table S3). In summary, these data revealed proteome-wide normalization of dysregulated proteins upon embryonic UBE3A reinstatement in the mICD mice and that the bi-allelic expression of 15q11-q13 genes does not impede this molecular rescue.

**Figure 4.**
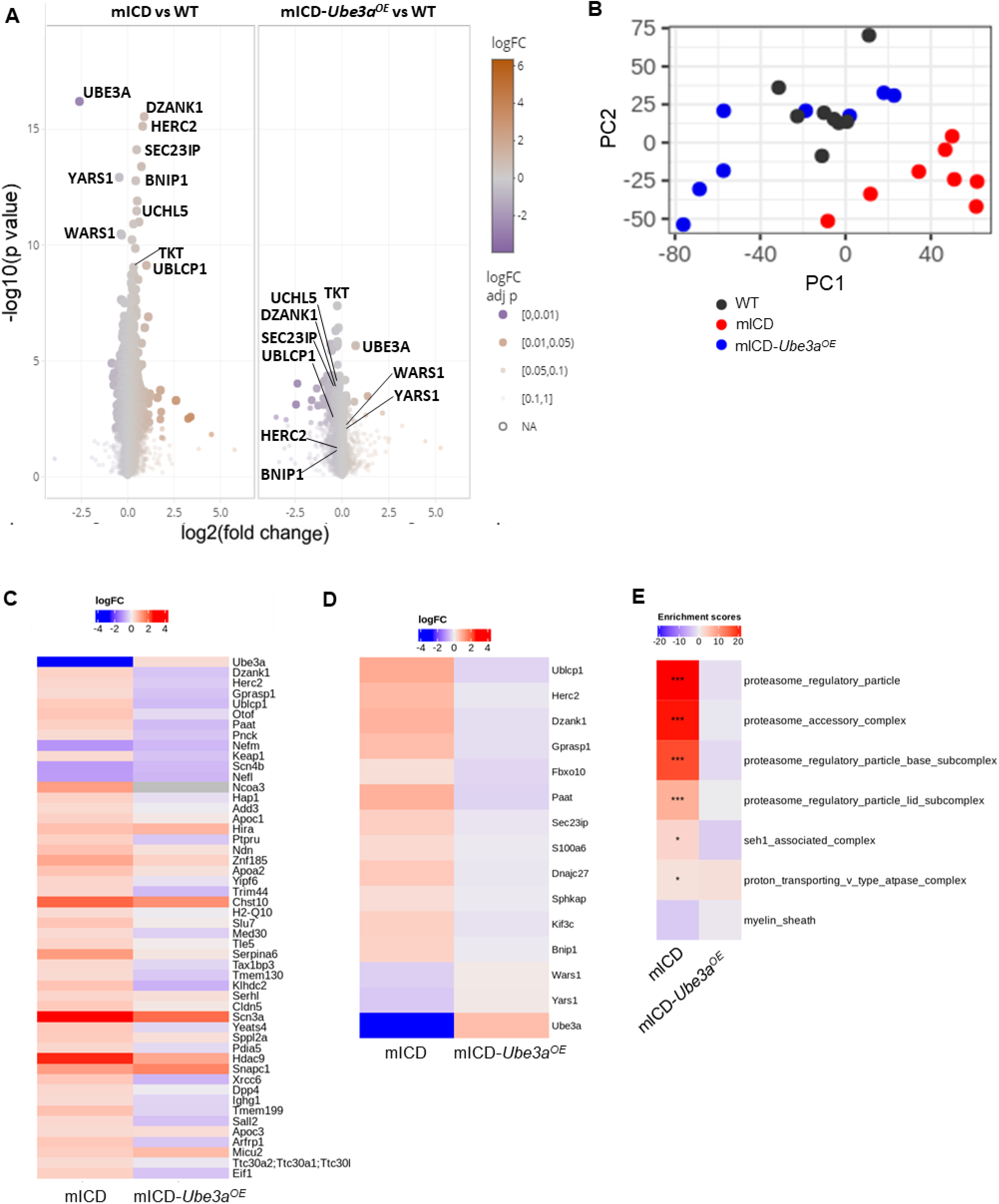
Embryonic transgenic UBE3A reinstatement in mICD mice rescues proteome alterations. (A) Volcano plots showing the magnitude of effect (log2 fold change on the x-axis) against the statistical significance (-log10 FDR-adjusted *p* values on the y-axis) for the data independent acquisition (DIA)-based proteomic profiling of adult mICD-*Ube3a^OE^* and mICD cortex (with respect to their WT control). (B) Principal component analysis (PCA) of all proteins shows a clear separation based on genotype (PC 2) and condition (PC 1). (C) Heatmap showing the top 50 differentially abundant proteins in the mICD cortex and its reversal in the mICD- *Ube3a^OE^* cortex when compared to the WT group (red: most abundant; blue: least abundant). Proteins are ordered by significance, with most significant at the top. Fold changes in mICD and mICD- *Ube3a^OE^* are relative to WT (D) Heatmap of the 15 overlapping proteins between the mICD and *Ube3a KO* mouse models (with respect to their WT controls). (E) Significantly enriched gene sets from the Gene Ontology - Cellular Component (GO:CC) are shown for the contrasts mICD vs WT and mICD-*Ube3a^OE^* vs WT (*** Benjamini Hoch FDR<0.001, **FDR<0.01, *FDR<0.05).

### Neonatal ASO-mediated UBE3A reinstatement in mICD mice partially rescues certain behaviors

Encouraged by the positive outcomes achieved through transgenic embryonic *Ube3a* overexpression in the mICD mice, our subsequent investigation aimed to assess whether treatment with a *Ube3a-ATS* - targeting ASO (hereafter referred to as tASO) of mICD mice would yield similar results to those seen in *Ube3*a KO-tASO treated mice. Consistent with our previous study, we administered 22 μg of tASO into the lateral ventricle of mICD or WT female mice (10). We also treated mice with a non-targeting scrambled (SCR) ASO control. This SCR was based on previously published non-targeting ASO sequences that demonstrated an absence of acute toxic manifestations after ICV injection in adult mice. This SCR also has similar nucleotide content and chemical modifications to the tASO (27). We first evaluated if the tASO would lead to bi-allelic activation of *Ube3a* and increased UBE3A protein expression. Following ASO injection, the cortex was extracted at two time points: P10 and P42 The first time point was chosen due to the anticipated peak effect of tASO on RNA and protein expression of UBE3A, and the second was chosen because it was the time point when mice enter the behavioral battery. The cortices of tASO and SCR treated mice were analyzed for RNA/protein expression, similar to the analysis performed in tASO-treated *Ube3a KO* mice (10) (Figure 5A). In mICD mice, the tASO led to highly efficient knockdown of *Ube3a-ATS* at P10 compared to SCR (Figure 5B), and *Ube3a* mRNA was expressed at a level higher than in WT (1.7x), likely reflecting the bi-allelic activation of *Ube3a* (Figure 5C). However, the increase in UBE3A protein level was comparatively modest (1.3x compared to WT) (Figure 5D). Similar results were observed at P42, which shows the sustained expression of UBE3A even at 6 weeks post-injection (Figure 5 C-D).

**Figure 5.**
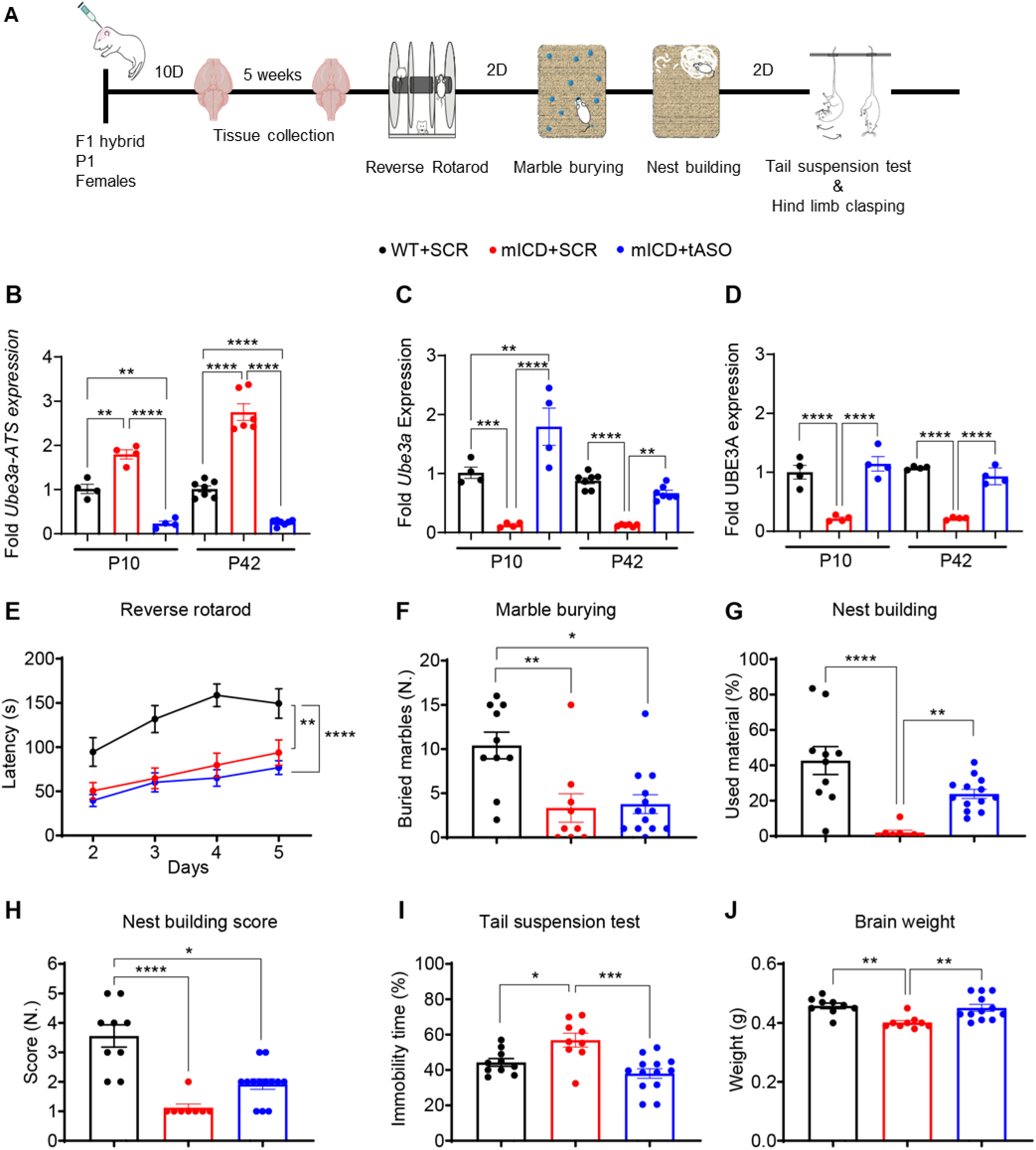
Neonatal ASO-mediated UBE3A reinstatement in mICD mice partially rescues certain behaviors. (A) Illustration of the timeline for execution of the behavioral battery and tissue collection. (B-C) Quantitative RT-PCR of cortex lysates of P1 injected WT+SCR (n=4), mICD+SCR (n=4) and mICD+ASO (n=4) collected at P10, WT+SCR (n=7), mICD+SCR (n=6) and mICD+ASO (n=7) collected at P42, analyzing *Ube3a-ATS* RNA (B) and *Ube3a* mRNA (C). (D) Capillary Western blot of the respective groups at P10 and P42 (n=4 per group). (E-I) Behavioral tests performed with WT+SCR (black), mICD+SCR (red) and mICD+tASO (blue) mice in the reverse rotarod (n=10; 9; 13), marble burying (n=10; 9; 13), nest building (n=10; 8; 13), quality of built nest (n=9; 8; 13), tail suspension (n=10; 8; 13). (J) Brain weight (n=9; 9; 12) was measured at the end of the battery. Data are represented as means ± SEM. *P* values are displayed as asterisks in the figure: not shown if *P* > 0.05, \**P* ≤ 0.05, \*\**P* ≤ 0.01, \*\*\**P* ≤ 0.001, \*\*\*\**P* ≤ 0.0001.

Akin to our prior ASO mouse study in *Ube3a KO* mice (10), six weeks post ASO injection, we conducted a compressed version of the behavioral battery to ensure that mICD-tASO treated mice would optimally express UBE3A during testing. Hence, mice were immediately subjected to the nest-building task after the marble burying test, bypassing the typical week-long acclimatization period to single housing. tASO treatment resulted in a significant improvement in the nest building test and a rescue of immobility time in the tail suspension test, as well as brain weight (Figure 5 G-J). However, no improvements were evident in the reverse rotarod or marble burying tests (Figure 5 E and F). Notably, we again did not detect a hind limb clasping phenotype in this cohort of mice (Figure S4A). Regarding body weight, we did not observe obesity in mICD compared to WT-SCR treated mice (Figure S4B). Additionally, mICD mice treated with tASO exhibited a similar trend in body weight measurements compared to the other two groups.

## Discussion

Angelman Syndrome is a debilitating neurological disorder caused by neuronal loss of function from the E3 ubiquitin ligase UBE3A. The scientific community has primarily concentrated on unraveling the function of UBE3A and its role in the pathophysiology of Angelman Syndrome (AS) and disease-modifying treatments attempt to increase neuronal UBE3A expression. In the AS genetic subtypes mICD and UPD, collectively representing approximately 20% of AS individuals, in addition to loss of UBE3A in neurons, bi-allelic expression of several imprinted genes within the 15q11-q13 region occurs. These genes are typically expressed only from the paternal allele. This cluster comprises the *MKRN3*, *NDN*, *MAGEL2*, and *SNHG14* genes, the loss of function of which contribute to Prader-Willi and Schaaf-Yang syndromes (28–31). However, it is not fully known whether their increased expression alone or in combination with the loss of *UBE3A* contribute to pathological consequences in the mICD and UPD genetic subtypes of AS.

Most mouse models of AS are *Ube3a*-centric and do not address the contribution of other genes affected in the 15q11-q13 region to AS pathophysiology. To investigate whether this cluster of genes within the 15q11-q13 region co-contribute to the behavioral defects observed in mICD and UPD patients, we employed an mICD mouse model (19) exhibiting bi-allelic expression of the *Mkrn3-Snhg14* gene cluster and *Ube3a* silencing, as validated by RT-PCR. We show that the mICD mouse model displays all core behavioral features previously observed in *Ube3a KO* mouse models of AS, further validating the robustness of this behavioral battery as a tool to study AS in mice with different genetic etiologies (20). Notably, the distance traveled in the open field arena was the only phenotype we could not reproduce in the mICD mouse model. This observation aligns with a meta-analysis study which showed that this test has the weakest statistical power (20). Notably, our initial analysis of the mICD model showed a phenotype in the tail suspension and hind limb clasping tests, which have also been observed in the *Ube3a KO* model (11). However, the hindlimb clasping phenotype could not be reproduced in subsequent mICD cohorts, suggesting this is not a strong phenotype. In addition, the tail suspension phenotype was lost upon crossing the mICD line with the *Ube3a^OE^* line indicating that this phenotype is either weak or sensitive to subtle changes in genetic background.

Comprehensive molecular analysis in mICD mice compared to WT controls revealed genotype-driven transcriptomic changes, with increased expression of transcripts mapping to the 15q11-q13 region, without enrichment for specific pathways. Furthermore, consistent with a previous proteomic analysis of *Ube3a KO* mice, the present study showed similar molecular alterations in both mICD and *Ube3a KO* cortex, with the proteasome pathway being significantly deregulated (26). Interestingly, some tendency of broader protein deregulation was observed in mICD compared to *Ube3a KO* mice. Perhaps, the increased expression of other 15q11-q13 genes in mICD mice is driving secondary/tertiary changes in the proteome, which otherwise are less prominent in the *Ube3a KO* mice. Taken together, our molecular and behavioral results imply that loss of *Ube3a* primarily drives molecular and behavioral changes in adult mICD mice.

Current therapeutic approaches focus on UBE3A reinstatement by ASOs targeting the *UBE3A- ATS* RNA that silences the paternally expressed *UBE3A* gene. However, it is possible that the bi-allelic expression of the *UBE3A-ATS* RNA in mICD/UPD patients would result in UBE3A overdosing upon ASO treatment. Given that clinical data may suggest that increased expression of UBE3A by itself or through its interaction with biallelically expressed genes in the 15q11-q13 locus could lead to pathological effects (reviewed by (17)), we tested the effect of expressing two *Ube3a* gene copies in the mICD mouse model. To this end, we mated mICD mice with *Ube3a^OE^* mice harboring two copies of the entire *Ube3a* gene on chromosome 3. In these double mutants, we observed a 1.5-2 fold increase in UBE3A protein levels, compared to WT. Behavioral testing of the mICD-*Ube3a^OE^* model demonstrated that *Ube3a* transgenic over-expression fully rescued all behavioral phenotypes observed in mICD mice. This suggests that in the selected tests, the phenotype is solely driven by the loss of UBE3A and the biallelic expression of paternal genes does not impact these phenotypes. Second, these results imply that there is no synergistic (pathogenic) interaction between bi-allelic expression of paternal genes in combination with UBE3A overexpression. Interestingly, prior to the initiation of the battery, mICD-*Ube3a^OE^* mice exhibited a body weight comparable to that of WT mice, indicating a rescue of the obesity phenotype observed in female mICD mice. However, over the course of the behavioral battery, we noted a progressive increase in body weight in mICD-*Ube3a^OE^* mice, unlike WT and mICD. Since our previous data did not show an effect of UBE3A over-expression on weight (18), this observation is puzzling and needs to be replicated. Proteomics analysis of cortical tissue of mICD-*Ube3a^OE^* mice further corroborated our findings indicating that *Ube3a* reinstatement could rescue the dysregulated expression of several top hits identified in the mICD cortex and also the proteasome related pathways, thereby normalizing global proteome comparable to WT cortex. Taken together, these experiments suggest that overexpression of *Ube3a* in the mICD background is well tolerated by the developing brain, consistent with previous reports in a *Ube3a^OE^* mouse model (18).

Encouraged by the rescue data of mICD-*Ube3a^OE^* double mutants, we next investigated the outcome of neonatal ASO application in the mICD mice. In mICD mice, *Ube3a-ATS* silences *Ube3a* on both alleles, therefore an ASO treatment results in the activation of both *Ube3a* alleles, and thus overall increased expression of UBE3A compared to *Ube3a KO* but also WT mice. At the molecular level, tASO treatment led to a pronounced silencing of the antisense transcript in mICD mice, indicating effective ASO targeting of both copies of the Ube3a-ATS. While a nearly two-fold increase of *Ube3a* mRNA above WT levels could be observed at P10, the increase for UBE3A protein is less pronounced (1.3x compared to WT). This is in line with previous UBE3A over-expression studies that showed that increasing *Ube3a* mRNA levels does not perfectly scale with UBE3A protein levels (18).

The robust molecular efficacy of tASO in UBE3A reinstatement was sustained for at least six weeks after treatment when the behavioral battery was started. However, only partial improvement in AS behavioral phenotypes was achieved. A complete rescue of immobility time in the tail suspension test and microcephaly was observed. Furthermore, improvements in the nest building task, both quantitatively and qualitatively, were noted. Conversely, no discernible effects were observed on marble burying and reverse rotarod phenotypes. These findings were unexpected, as we showed previously that treatment with the same tASO in *Ube3a KO* mice did not yield improvement in the nest building task, but improved performance on the reverse rotarod task. The window for effective intervention in the reverse rotarod task closes after P21, suggesting that the lack of rescue in mICD for this phenotype is not likely attributed to timing of treatment (32). Possibly, the absence of UBE3A during embryonic development, coupled with dysregulation of the duplicated genes, might affect the formation of neuronal networks crucial for motor coordination assessed in the reverse rotarod task. Conversely, a lack of improvement in the nest building task following ASO treatment of *Ube3a KO* mice could be attributed to the lower levels of UBE3A reinstatement achieved in that line, which was approximately 70% relative to WT at P7 (10). Further (replication) studies are needed to understand these differences in behavioral rescue obtained by ASO treatment.

In conclusion, our results demonstrate that bi-allelically expressed genes within the 15q11-q13 region have a minimal contribution to AS pathology in mICD mice and that embryonic, transgenic *Ube3a* reinstatement can effectively rescue molecular and behavioral deficits in mICD mice. In addition, application of *Ube3a-ATS* targeting ASO in mICD mice rescues several phenotypes. Thus, treatment options aiming at re-activating paternal *Ube3a* expression, such as *Ube3a-ATS* knockdown via antisense oligonucleotide therapy currently under investigation in Phase I clinical trials for AS, should be considered as potentially safe and effective options for ICD and UPD patients.

## Material and Methods

### Mouse husbandry and breeding

The mICD mouse line harboring a floxed rabbit ß-globin transcriptional terminator between U1 and the PWS-IC was obtained from Prof. James Resnick, University of Florida (19). For behavioral experiments, female C57BL/6J mICD were crossed with male 129S2 mice to generate mICD mice and their WT littermates in an F1 hybrid 129S2-C57BL/6J background. Mice were genotyped during weaning, three to four weeks after birth. Two weeks before the behavioral battery, female mICD and WT mice were shipped from the animal facility of Charles River Laboratories, Sulzfeld, Germany to that of the Erasmus MC (EMC), where all behavioral experiments were conducted. Given that we observed some aggressive behavior among males bred at our external facility, only adult female mICD mice and control littermates in the F1 hybrid 129S2-C57BL/6J background were used for the initial behavioral battery and following molecular analyses (qPCR and WB).

For behavioral experiments with miCD-*Ube3a^OE^* mice, C57BL/6J mICD females were crossed with 129sv *Tg-Ube3a-FL* OE2x (*Ube3a^OE^*) males at Erasmus MC facilities, yielding F1 hybrid 129S2-C57BL/6J miCD-*Ube3a^OE^*, WT and mICD littermates. Both males and females were used for behavioral and follow-up molecular analyses (qPCR and WB).

For behavioral experiments with mICD and WT animals injected with the tASO or SCR, pregnant dams (C57BL/6J mICD) were shipped at E15 from the animal facility of Charles River Laboratories, Sulzfeld, Germany to that of the EMC, where all behavioral experiments were conducted. All pregnant dams were single housed until delivery.

Mice born at the EMC were genotyped 4–7 days after birth and all mice were re-genotyped at the end of each experiment. Animals were housed in individually ventilated cages (IVC; 1145T cages from Techniplast) in a barrier facility with a temperature of 22 ± 2°C, a 12:12-hour light: dark cycle, and provided with ad libitum food [801727CRM(P) from Special Dietary Service] and water. Mice were group- housed, with 2 to 4 animals of the same sex per cage, until a week before the nest-building task, after which they were single-housed.

### ICV injection of newborn mice

The tASO or SCR were prepared by diluting them to a final concentration of 22 μg/μL in PBS supplemented with 0.3% Fast Green. The injection procedure closely followed the method outlined by Milazzo et al. 2021. In brief, P1 mice were cryo-anesthetized and injected using a glass pipette with a tip diameter of 0.5–0.7μm, attached to a 25μL syringe (Hamilton, model 1702N). Mice were injected in the lateral ventricle by positioning the needle at the midpoint between the right eye and the lambda intersection of the skull and then lowering it to a depth of approximately 3 mm. A volume of 1μL of either tASO, SCR, or PBS was administered at a flow rate of 0.5 μL/min using a CMA 400 Syringe Pump (Harvard Apparatus).

The tASO utilized in this study is identical to the one described in Milazzo et al. 2021, designated as ASO RTR26266, with the following nucleotide sequence: TCCaacttaataaCCT. The selection of the SCR ASO, RTR22946, was based on a list of non-targeting ASOs published by (27). Its nucleotide sequence is: CcAAAtcttataataACtAC. In both sequences, capital letters represent PS-LNA modifications, where all LNA- C nucleotides incorporate the 5-methyl cytosine modification, while lowercase letters represent PS-DNA.

### Behavioral testing

Mice were randomly assigned to experimental groups, while taking a male/female balance into account. The weight of the animals was measured on the same day as the start of the behavioral battery, before the first test and on the day of euthanasia. Between 9 and 12 weeks of age, mICD, mICD-*Ube3a^OE^* and WT mice were subjected to a standardized behavioral battery performed by a researcher blind to the genotype, during the light period of the light/dark cycle. No animals were excluded. Data from a previously published meta-analysis was used to calculate the sample size for the behavioral experiment (20). Behavioral tests with mice not ICV injected were performed in the following order: 5 consecutive days of reverse rotarod; two days of break; open field (not performed with mICD-*Ube3a^OE^* and ASO-treated cohorts); two days of break; marble burying; a week of pause to allow adaptation to single-housing; 5 days of nest building (weight of the nest was assessed at the same hour every day in the experiment assessing differences between mICD and WT animals, and only on day five in the experiment assessing differences between mICD- *Ube3a^OE^*, mICD and WT animals); two days of break; forced swim test (only performed in experiment assessing differences between mICD-*Ube3a^OE^*, mICD and WT animals); two days of pause; tail suspension test. The behavioral tests were performed in the following sequence with 6-week-old female mice that received ICV injection at birth with either tASO, SCR, or PBS: reverse rotarod for 5 days, 2-day break; marble burying, on the same day mice were single-housed and the nest building test commenced, continuing for 5 days; 2-day break; tail suspension test. All battery tests, with the exception of the tail suspension test, were performed as previously described (10, 20). For the tail suspension test mice were suspended in a frame, 40cm above the base, using a paper adhesive tape placed approximately 1 cm from the tip of the tail, and videotaped for 5 min. The immobility time was measured by scoring manually (stopwatch) for 5 minutes the amount of time (seconds) the mouse was not moving. The first and last 10 seconds of the recording were scored on a scale from 0 to 3 based on hind-limb clasping and the average score was used for hind-limb clasping analysis. The scale was adapted from a previous study: 0 = both hindlimbs were placed outward away from the abdomen, 1 = one hindlimb was retracted or both hindlimbs were partially retracted toward the abdomen without touching, 2 = both hindlimbs were partially retracted toward the abdomen and were touching the abdomen without touching each other, 3 = both hindlimbs were fully clasped and touching the abdomen (33). Additionally, at the end of the nest building task, photographs of the nest were taken for following nest quality analysis. The nests were rated on a scale from 0 to 4 based on what was reported in a previous study (8), in summary if no nesting material was used a score of 0 was assigned, if a minimal quantity of nesting material was used to build a rudimental nest a score of 1 was given, if more nesting material was used to build a disorganized nest a score of 2 was assigned, if most nesting material was used to build an organized nest with defined borders a score of 3 was given, and if >90% of nesting material was used to build a well-defined circle-like structure a score of 4 was assigned.

### Tissue processing and RNA/protein extraction

Mice were euthanized by cervical dislocation under anesthesia with isoflurane. The cortex, hippocampus and striatum were extracted immediately after by snap freezing and stored at -70°C till further processing. For qRT-PCR, RNA sequencing and Capillary western blotting, tissues were homogenized in tubes prefilled with ceramic beads (MagNa Lyser Green Beads, Roche, # 03358941001) using MagNA Lyser instrument (Roche). Subsequently, the lysate was processed using a spin-column based kit (Norgen Biotek Corp., #47700) for sequential extraction of DNA, RNA and Protein, according to the manufacturer’s instructions.

### Real-time qRT-PCR

Total RNA quantity and quality was determined using the NanoDrop 1000 spectrophotometer (DeNovix, Life Science Technologies). From this, 50ng RNA was first treated with the TURBO DNase enzyme (Thermo Fisher, #AM2238) according to manufacturer’s directions, to remove any residual DNA contaminants and diluted to 2ng/ul. Subsequently, each RNA sample was reverse-transcribed and quantitative real-time PCR was performed simultaneously in a single step using iTaq Universal SYBR Green One-step kit (Bio-Rad, #1725150), in the Lightcycler instrument (Roche). Primers were obtained from Microsynth AG and sequences are listed in Supplemental Table 1. Each sample were measured in triplicates. Using the mean Ct-value of these triplicates, real-time qRT-PCR data were analyzed by normalizing each transcript to GAPDH level, with each value represented as a fold change with respect to the control group.

### Capillary western blot

Protein levels of UBE3A in the mouse brain were analyzed by automated capillary western blotting (Sally Sue, Protein Simple). All experimental steps were carried out according to the manufacturer’s instructions. Briefly, after protein extraction and quantification by BCA protein assay (ThermoFisher, #23225), a final sample concentration of 0.5 mg/ml was loaded to the capillary cartridges (12–230 kDa Peggy Sue or Sally Sue Separation Module, #SM-S001). Chemiluminescent protein detection was performed using the Anti Rabbit Detection Module (#DM-001) and the area under the chemiluminescent curve for each protein probed was obtained from the Compass for SW software (Version 4.1.0, Protein Simple). The values for UBE3A were then normalized to that of HPRT protein, and represented as a fold change with respect to the control group. Antibodies used are UBE3A (Bethyl, #A300-352A, 1:50), HPRT (Abcam, #ab109021).

### Immunoblotting

Twenty micrograms of protein lysate, extracted from one cerebral hemisphere, were separated on a precast 4–12% Criterion XT Bis-Tris gel (Bio-Rad) and transferred to a nitrocellulose membrane using TurboBlot (Bio- Rad). After blocking for 1h, at room temperature, with blocking buffer, the following antibodies were used: anti-UBE3A (mouse, anti-E6AP, MilliporeSigma, # E8655, 1:1000) and anti-GAPDH (rabbit, Cell Signaling, # 2118, 1:1000). The next day, the membranes were probed with secondary goat anti-mouse antibody (LI-COR Biosciences, IRDye 800CW, # 926-32210, 1:15,000) and goat-anti-rabbit antibody (LI-COR Biosciences, IRDye 680RD, # 926-32221, 1:15,000). The membranes were scanned using Odyssey CLx (LI-COR Biosciences) and quantified using the Odyssey 3.0 software.

### RNAseq and data analysis

Total RNA was isolated from seven mICD and seven WT adult female cortex. An RNA integrity of above 8 for all samples was confirmed post extraction using the Tapestation RNA ScreenTape from Agilent. 500 ng of total RNAs were used to prepare sequencing libraries using the TruSeq Stranded total RNAs Library Prep Kit (Illumina) according to the manufacturer’s instructions. All libraries were multiplexed into a single pool which was sequenced across 2 lanes of a SP flow cell of an Illumina NovaSeq 6000 instrument at the Genomics 360 Labs, Roche. On average, each library generated approximately 100 million paired-end 75 bp reads.

Base calling was performed with BCL to FASTQ file converter bcl2fastq2 version 2.20.0 (BCL to FASTQ file converter, available online at Illumina Inc.). FASTQ files were quality checked with FastQC version 0.11.9 (available online at Babraham Institute: https://www.bioinformatics.babraham.ac.uk/projects/fastqc/). Paired-end reads were mapped onto the mouse genome (build mm39) with read aligner STAR version 2.7.10b using default mapping parameters (34). Alignment metrics were determined with Picard version 2.25.1 (Picard Toolkit, 2018. Broad Institute, GitHub Repository: http://broadinstitute.github.io/picard/; Broad Institute). Read sequences and alignments were quality checked with MultiQC version 1.12 (35). Numbers of mapped reads for all RefSeq and/or Ensembl transcript variants of a gene were combined into a single value (i. e. read count) assuming reverse stranded library by featureCounts version 2.0.1 (36) and normalized as tpm (transcripts per million). We applied the edgeR algorithm (37) for differential gene expression analysis. For gene set enrichment analysis, we performed the adapted CAMERA algorithm implemented in the ribiosNGS package (Next-generation sequencing data analysis with ribios. https://github.com/bedapub/ribiosNGS).

### Proteomics and data analysis

Total protein profiling of adult mouse cortex tissue from WT vs mICD and WT vs *Ube3a KO*) was performed at Biognosys AG (Schlieren, Switzerland) and the proteomic profiling of adult mouse cortex tissue from WT vs mICD vs mICD-*Ube3a^OE^* was performed at Proteomics 360 Labs at Roche using Hyper Reaction Monitoring (HRM™) label-free discovery proteomics workflow. All measurements were performed in a randomized, blinded fashion and balanced for genotype and age (4 males, 4 females per genotype). Sample preparation, Data-independent acquisition (DIA) mass spectrometry and data analysis were performed as described in Pandya et al 2022 and described here.

Tissue samples were denatured using Biognosys’ Denature Buffer, and reduced and alkylated using Biognosys’ Reduction and Alkylation Solution for 60 min at 37°C. Subsequently, digestion to peptides was carried out using trypsin (w/w ratio 1:50 Promega) overnight at 37°C. Peptides were desalted using an Oasis HLB µElution plate (Waters) according to the manufacturer’s instructions and dried down using a SpeedVac system. Peptides were resuspended in LC solvent A (1 % acetonitrile, 0.1 % formic acid (FA)) spiked with Biognosys’ iRT kit calibration peptides. Peptide concentrations were determined using a UV/VIS Spectrometer (SPECTROstar Nano, BMG Labtech) and 3.5 µg of peptides were injected to an in-house packed reversed phase column on a Thermo Scientific™ EASY-nLC™ 1200 nano-liquid chromatography system connected to a Thermo Scientific™ Orbitrap™ Exploris 480™ mass spectrometer equipped with a Nanospray Flex™ ion source and a FAIMS Pro™ ion mobility device (Thermo Scientific™). LC solvents were A: water with 0.1 % FA; B: 80 % acetonitrile, 0.1 % FA in water. The nonlinear LC gradient was 1 – 50 % solvent B in 210 minutes followed by a column washing step in 90 % B for 10 minutes, and a final equilibration step of 1 % B for 8 minutes at 60 °C with a flow rate set to 250 nL/min. The FAIMS-DIA method consisted per applied compensation voltage of one full range MS1 scan and 34 DIA segments as adapted by (38, 39). Total protein profiling was performed using Biognosys’ Hyper Reaction Monitoring (HRM™) label-free discovery proteomics workflow. The mass spectrometric data were analyzed using Pulsar search engine as implemented in Spectronaut software (version 16.2, Biognosys), the false discovery rate on peptide and protein level was set to 1 %. A mouse UniProt .fasta database (Mus musculus, 2022-07-01, 17’125 entries) was used for the search engine, allowing for 2 missed cleavages and variable modifications (N-term acetylation and methionine oxidation). The generated directDIA library was used for further analysis and the HRM measurements analyzed with Spectronaut were normalized using global normalization on the median.

For each dataset, relative abundance measurements were log2 transformed and Gene Ontology (GO) annotations for each protein were downloaded from UniProt (40). Principal Component Analysis (PCA) was performed using the whole proteomics dataset. PCA was used to visualize the data in a reduced space capturing the highest variability, to potentially highlight the differences between the experimental conditions or due to covariates. Outliers were detected using the robust Mahalanobis distance computed from the first 3 principal components.

Differential abundance analysis between conditions was based upon linear models applied to each protein independently using the limma package in R (41). Specific contrasts of interest were defined in each dataset. For both datasets, no covariate was identified as impacting the data, leading to the following linear models: First dataset: y ∼ Group*Mouse_Model, with Group referring to KO or WT, Mouse_Model referring to *Ube3a KO* or mICD animal models. Second dataset: y ∼ Group, with Group referring to WT, mICD or mICD-*Ube3a^OE^* groups. Moderated t-tests were then performed using the Empirical Bayes method, which moderates the variance (42). P-values were adjusted for multiple comparisons using the Benjamini– Hochberg method to control the false discovery rate (FDR). Proteins with an adjusted *p*-value below 0.05 were considered differentially expressed. For gene-set enrichment analysis (GSEA), we considered Gene Ontology (GO) collections related to Biological Processes, Molecular Functions and Cellular Components. We report here the results from the GO:CC collection (43). For each contrast of interest, all genes/proteins were ranked and compared to the sets of genes from the database. A normalized enrichment score was then computed for each gene set using the camera function in the limma package, which considers the potential correlation between genes. In our case, we considered a fixed inter-gene correlation at 0.01, to better rank biologically interpretable gene sets (44). The associated *p*-value estimates the statistical significance of the score, based on the null distribution of the Kolmogorov–Smirnov statistic (45). As multiple enrichment tests were performed, the Benjamini-Hochberg method was used to correct for multiple-hypothesis testing.

### Statistics

All statistical analyses were done using GraphPad Prism software (v7.0, GraphPad Software Inc.,) and P value less than 0.05 was considered significant. Western blot and qRT-PCR data were analyzed using t-test or a 1-way ANOVA, 2 sided, with genotype as independent variable, and followed by Tukey’s post hoc test. Before performing any statistical test on the behavioral data, the samples were analyzed for normal distribution. To analyze data from the open field, marble burying, tail suspension test, forced swim test, hind-limb clasping, nest quality and quantity of nest material used on day 5, brain and body weight we used a parametrical or non-parametrical 1-way ANOVA, 2 sided, with the variable genotype as independent variable, followed by Tukey’s post hoc test. Both the reverse rotarod and nest building, where quantity of material used was measured over five days, were analyzed with a 2-way repeated-measure ANOVA, 2 sided, followed by Tukey’s post hoc test, with genotype and time as independent variables. All data are plotted as mean ± SEM. *P-*values show the significance level for the measurement and are indicated as asterisks in the figures and not shown if *p*-value is more than 0.05. Detailed information about the statistical test used for each experimental measurement can be found in Supplemental Table 2.

### Study approval

All animal experiments were conducted in accordance with the European Commission Council Directive 2010/63/EU (CCD license AVD101002016791 and AVD10100202216352).

## Data availability

RNAseq data will be deposited in the NCBI’s Gene Expression Omnibus database and made available close to manuscript publication. All .raw files from proteomics experiments will be made available on Massive server close to manuscript publication.

## Author contributions

CM, SW and RA performed behavioral characterization, CM performed ASO injections. RN and CM performed RNA and protein quantification assays, and all statistical analysis. RN prepared samples for and coordinated proteomics and RNAseq experiments. SB supported proteomics data analysis. YE, TK, and SC conceived the study and supervised the study. EM also supervised the study. CM and RN made the figures and wrote the manuscript which was reviewed by all authors.

## Acknowledgements

mICD model mice were a generous gift from James Resnick, and *Ube3a^OE^* mice were a generous gift from Ben Philpot. Initial characterization of mICD mice and treatment with ASOs was funded by Roche. Characterization of the mICD mice crossed with *UBE3A^OE^* mice was funded by the Angelman Syndrome Foundation (ASF). CM was supported by grants from Associazione Angelman and FROM. RN was supported by funds from the Roche Postdoctoral Fellowship. We thank Manuel Tzouros and Sabrina Golling (Proteomics 360 Lab, Roche) for proteomics support, Kim Schneider and Kerstin Oelkers (Genomics 360 Lab, Roche) for RNAseq support. We thank Martin Ebeling and Roland Schmucki for RNAseq data analysis support (PS-PMDA, Roche).

