## Supplemental Figures for "UBE3A reinstatement restores behavior and proteome in an Angelman Syndrome mouse model of Imprinting Defects"

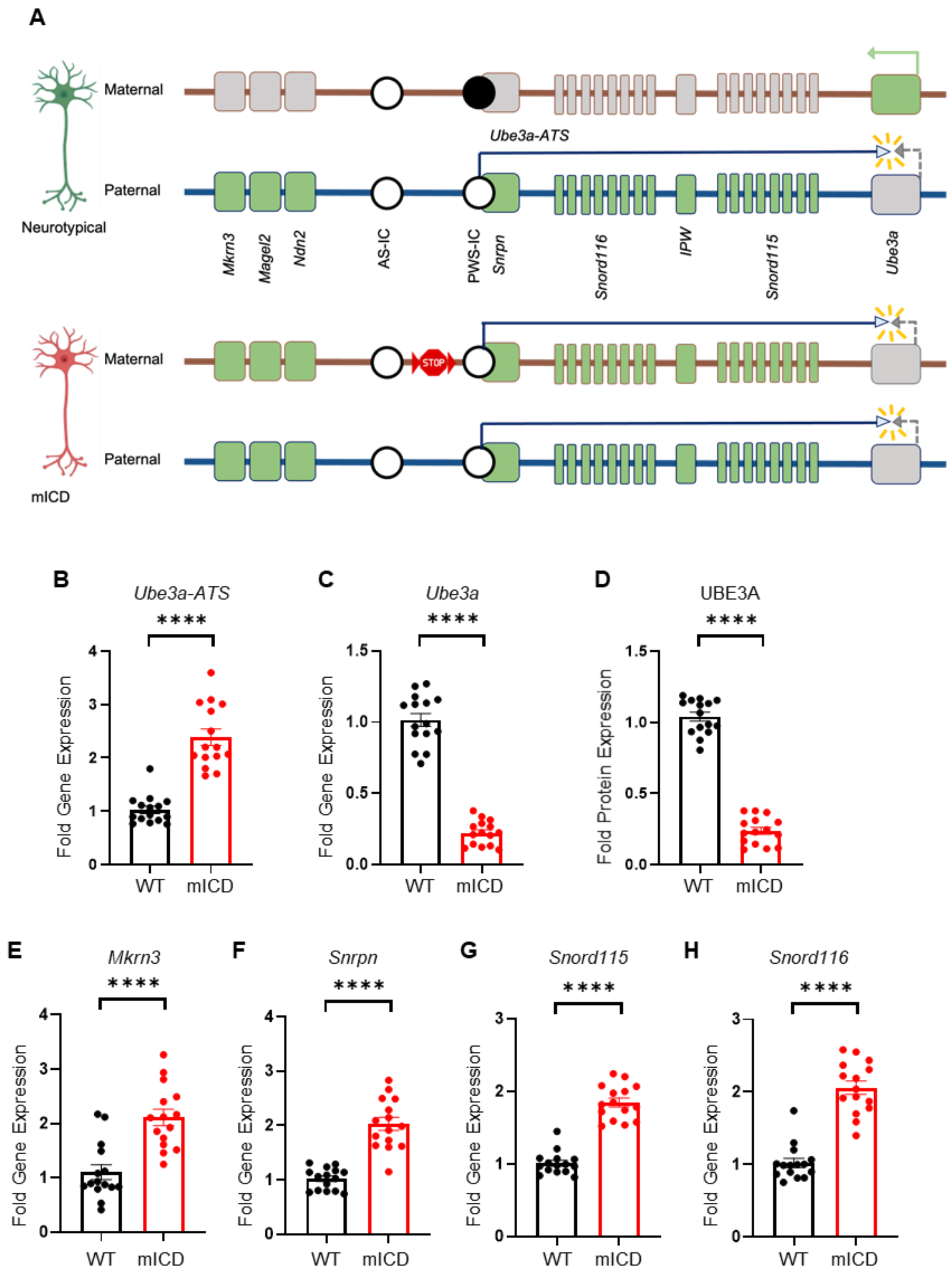


**Figure S1. Maternal imprinting defects lead to gene expression changes.** (A) Schematic representation of the 15q11-q13 locus on mouse chromosome 7 in control neurons (upper panel) and mICD neurons (bottom panel). Insertion of a transcription termination cassette (red stop sign) between the AS and PWS imprinting centers in the maternal allele, leads to activation of the paternally expressed genes: *Mkrn3, Magel2, Ndn* and *Snhg14*. The *Snhg14* transcription unit produces a polycistronic transcript comprising, among other transcripts, the *Snrpn* mRNA, the small nucleolar RNAs *Snord115* and *Snord 116*, the noncoding exons IPW, and the *Ube3a-ATS* long non-coding RNA. Transcription of the *Ube3a-ATS* results in silencing of *Ube3a*. Green and grey squares represent active and silenced genes respectively. Methylated and unmethylated imprinting centers are illustrated with filled and empty circles respectively. (B, C, E, F, G, H) Gene expression fold change in the cortex of mICD mice (n=15), relative to WT controls (n=15), measured in terms of RNA expression. (D) Capillary Western blot of cortex lysates of mICD (n=15) and WT mice (n=15), obtained 1 week after completion of the behavioral battery. In all graphs, data are represented as means ± SEM. *P* values, calculated via T-test, are displayed as asterisks in the figure: not shown if *P* > 0.05, **P* ≤ 0.05, ***P* ≤ 0.01, ****P* ≤ 0.001, *****P* ≤ 0.0001.

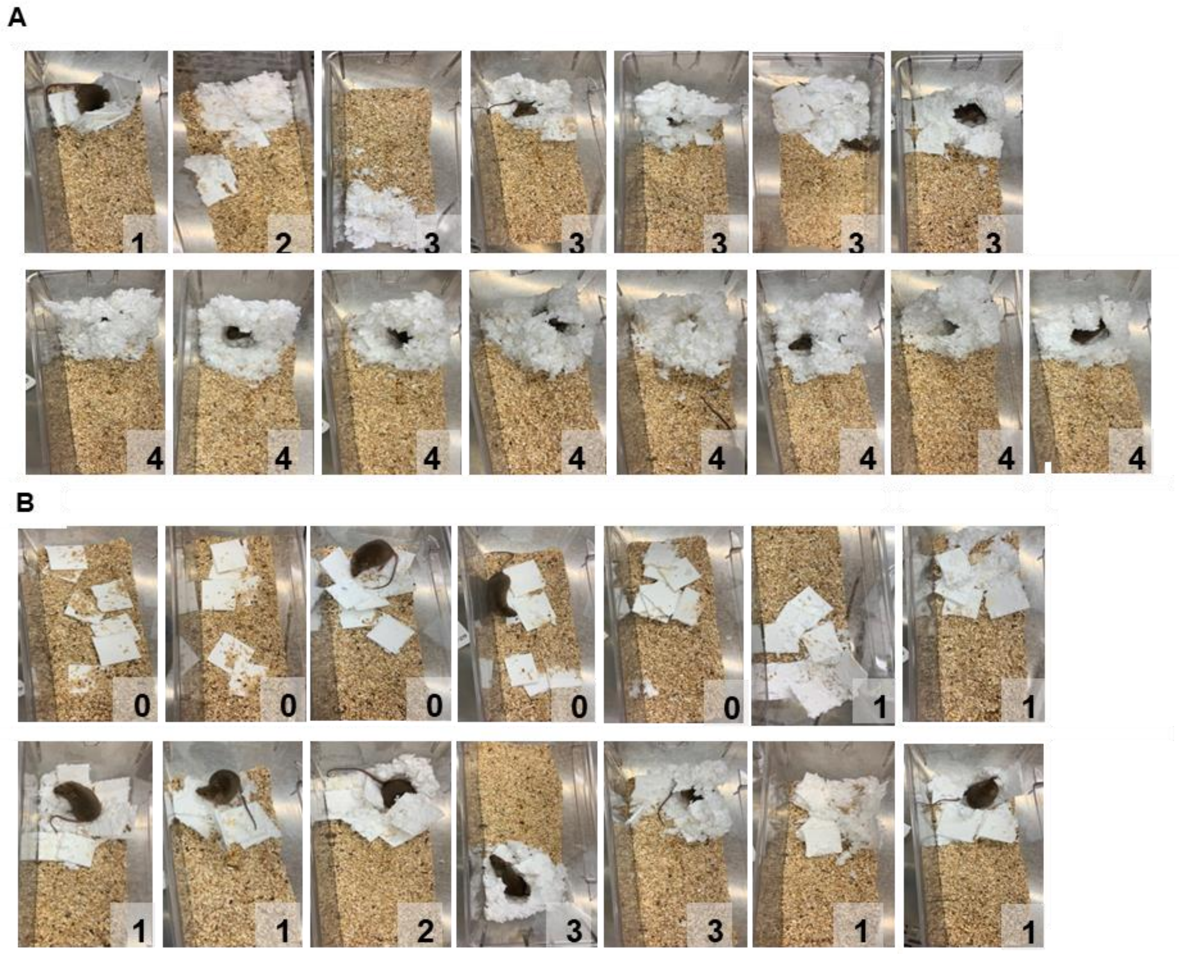


**Figure S2. mICD mice build nests of poorer quality compared to WT controls.** Images of nests built by WT mice (n=15), panel A, and miCD mice (n=14), panel B. The score for nest quality, on a scale from 0 to 4, is indicated at the bottom right corner of each image.

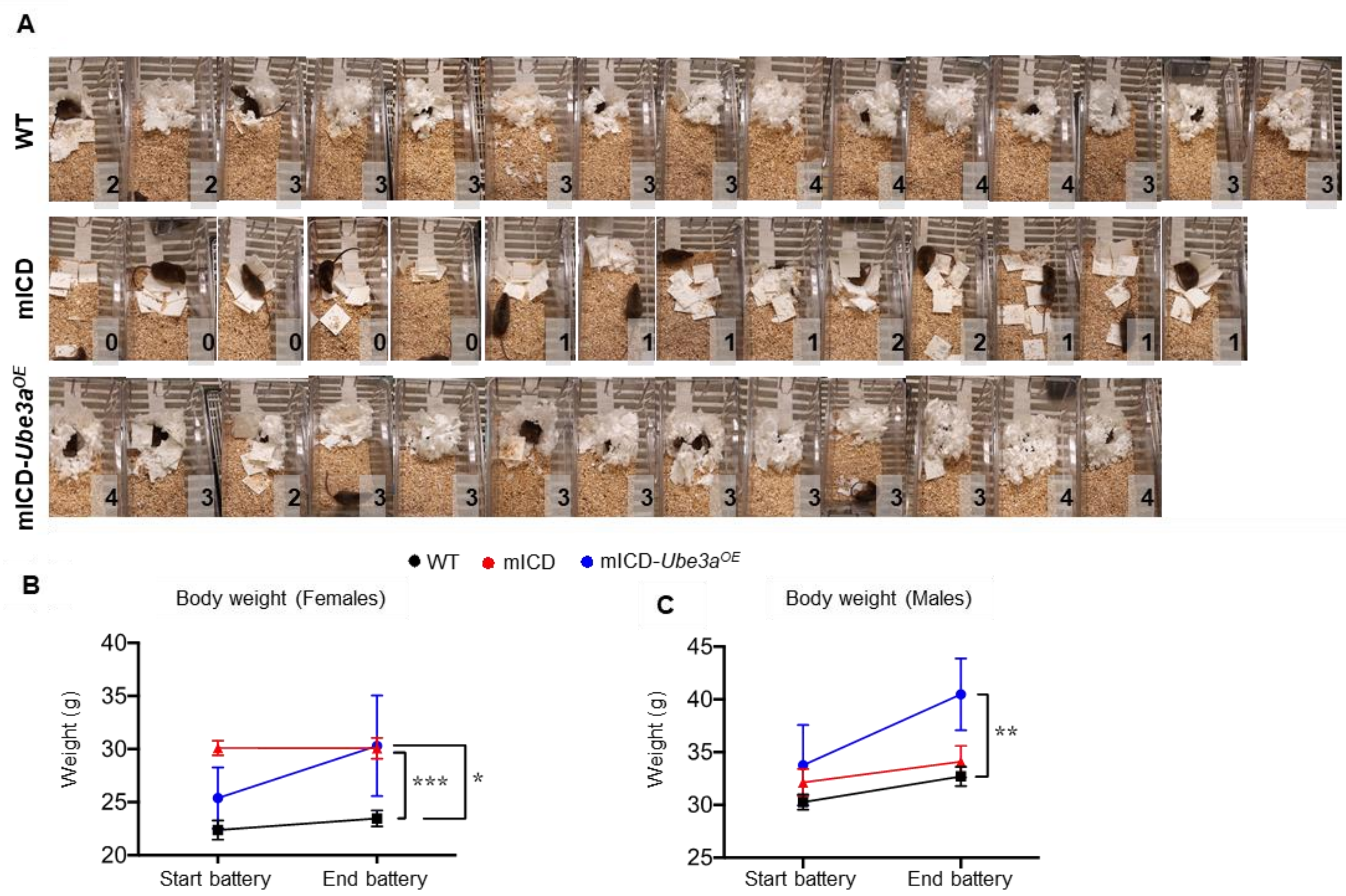


**Figure S3. mICD-*Ube3a*$^{OE}$ mice build nests of similar quality to WT controls and gain weight over time.** (A) Images of nests built by WT (n=15), mICD (n=14), and mICD-*Ube3a*$^{OE}$ mice (n=13). The score for nest quality, on a scale from 0 to 4, is indicated at the bottom right corner of each image. Body weight was measured in females (B) and males (C) WT (black; n= 6, 9); mICD (red; n = 7, 6) and mICD-*Ube3a*$^{OE}$ (blue; n= 8, 7) mice, the same day as the start of the battery, before the first test, and on the day of euthanasia. Two-way ANOVA with Tukey's post-hoc test used to assess changes in body weight.

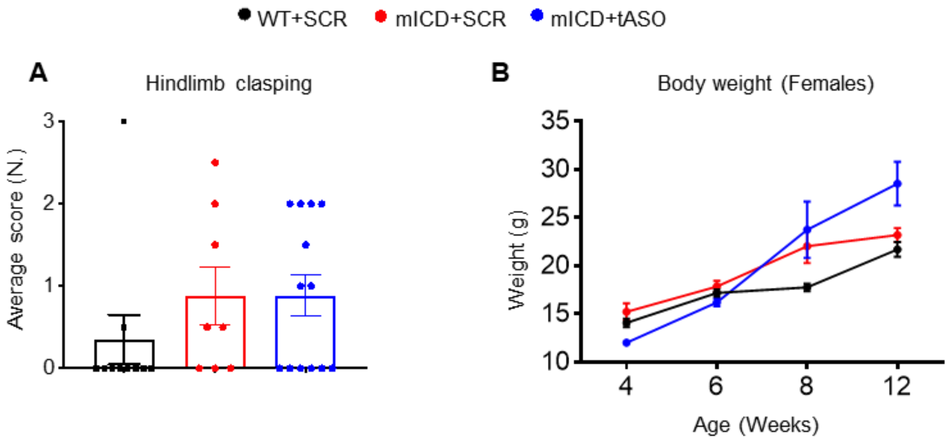


**Figure S4. Neonatal ASO-mediated UBE3A reinstatement in mICD mice partially rescues certain behaviors.** (A) Hind limb clasping test performed with WT-SCR (black, n= 10), mICD-SCR (red, n=8) and mICD-tASO (blue, n= 13). (B) Body weight was measured at different ages - WT-SCR 4 and 6 weeks (n=10), 8 weeks (n=5), 12 weeks (n= 4); mICD-SCR 4 and 6 weeks (n= 8), 8 weeks (n=4), 12 weeks (n= 3); mICD-tASO 4 and 6 weeks (n=13), 8 weeks (n=6), 12 weeks (n= 7). Data are represented as means ± SEM. *P* values are not shown if $P > 0.05$.
